# An expedient, biology-laboratory-compatible method for preparing functional perfluoropolyether fluorosurfactants for droplet microfluidics

**DOI:** 10.64898/2026.03.28.714914

**Authors:** Chase Akins, Jessica L. Johnson, Gyorgy Babnigg

## Abstract

Biocompatible fluorosurfactants are essential for many droplet microfluidic workflows but are often obtained from commercial sources because published syntheses of perfluoropolyether (PFPE)-based surfactants typically require acid chloride intermediates and chemistry-oriented purification methods. These requirements can limit access for biology and clinical laboratories seeking low-cost or customizable surfactant systems. Here we describe a practical method for preparing functional PFPE-based fluorosurfactant materials by direct carbodiimide coupling of functionalized PFPE carboxylic acids(Krytox 157 FSH) to amine-containing head groups under laboratory-accessible conditions. Using this approach, we prepared a PFPE-polyethylene-glycol (PFPE-PEG) material from Jeffamine ED900 and a PFPE-Tris material from Tris base.

Because these products were not fully structurally characterized, we present them as functional reaction products and evaluate them by use in biomicrofluidic workflows rather than by definitive compositional assignment. PFPE-Tris was useful for generating relatively uniform small droplets, whereas the PFPE-PEG preparation supported a broader range of biological applications. These materials were used in genomic library screening for β-glucosidase activity, thermocycling-associated droplet workflows, and protein crystallization experiments. In addition, the PFPE-PEG preparation improved emulsion behavior in many protein crystallization screens that were unstable with a commercial droplet oil used in our laboratory.

This method reduces the practical barrier to in-house fluorosurfactant preparation and allows biology-focused laboratories to explore head-group chemistry, oil composition, and operating conditions without complete reliance on commercial reagents. The results support this workflow as a useful entry point for biomicrofluidics laboratories, while also highlighting the need for careful interpretation of thermocycled droplet assays and for future analytical characterization of the resulting materials.

**Significance statement:** Droplet microfluidics relies on fluorosurfactants that are often costly and difficult to synthesize outside of chemistry-focused settings. We describe a simple, biology-laboratory-compatible approach for generating functional perfluoropolyether-based fluorosurfactant materials using direct carbodiimide coupling and straightforward cleanup. The resulting materials supported multiple biomicrofluidic workflows in our laboratory, including enzymatic screening and protein crystallization, and provide a practical route for groups seeking lower-cost and more customizable surfactant systems.

## Introduction

Droplet microfluidics is widely used for compartmentalized biochemical and biological assays, including enzyme screening, cellular analysis, nucleic acid amplification, and protein crystallization [1–3]. In many such systems, fluorinated oils are preferred as the continuous phase because they combine low flammability, chemical inertness toward many biomolecules, suitable viscosity for microchannel flow, and gas permeability across the droplet interface [1,2]. Stable operation, however, depends on fluorosurfactants that can support water-in-oil emulsions without strongly interfering with biological processes.

Many commercially available industrial fluorosurfactants are poorly suited to biological droplets. Their ionic character, interfacial behavior, or limited emulsion performance can reduce utility in biomicrofluidic assays. In contrast, PFPE-PEG-based surfactants derived from Krytox-type PFPE tails and hydrophilic amine-containing head groups are widely used because they often combine practical biocompatibility with useful thermostability [4–7]. A common formulation is the PFPE-PEG-PFPE triblock architecture often referred to informally as a Krytox-Jeffamine (KryJeffa) surfactant. Despite their importance, these materials are frequently purchased rather than synthesized in-house. Published synthetic routes often proceed through an acid chloride intermediate and may rely on reagents, purification workflows, or synthetic chemistry infrastructure that are not standard in biology or clinical laboratories [5,6]. This creates a cost and accessibility barrier, especially for groups that wish to test alternative head groups, different oils, or application-specific formulations.

We therefore explored whether functional PFPE-based fluorosurfactant materials could be prepared using direct carbodiimide coupling of Krytox 157 FSH to amine-containing head groups. Carbodiimide chemistry is familiar to many life-science laboratories, can be performed under mild conditions, and does not require acid chloride preparation. In the present work, we focused on reaction with Jeffamine ED900 to generate a PFPE-PEG material and with Tris base to generate a PFPE-Tris material. We then assessed these materials in several workflows relevant to biomicrofluidics.

Because the resulting products were not fully chemically characterized, we avoid assigning precise molecular structures or product distributions. Instead, we focus on practical function: droplet-generation behavior, compatibility with genomic library screening for β-glucosidase activity, behavior in thermocycling-associated workflows, and use in protein crystallization droplets

## Materials and Methods

### Reagents and materials

Miller-Stephenson Krytox 157 FSH was obtained from Grainger (Cat #35RR09). Jeffamine ED900 (Sigma, 14527) was used as the amine-containing PEG reagent, stored at 4°C. 1-Ethyl-3-(3-dimethylaminopropyl)carbodiimide (EDC, Sigma, E7750) was stored at −20°C and allowed to warm to room temperature for at least 3 h before use. Novec HFE-7500 fluorinated oil (Oakwood Chemical, 051243-1Kg) was used as the primary carrier oil unless otherwise noted. FC-40 (Sigma, F9755) was used in selected thermocycling experiments. PEG 35,000 flakes (Sigma, 81310) were used for cleanup. Additional reagents included MES monohydrate buffer (Sigma, 69892), Trizma base (Sigma, T1503), OptiPrep (Sigma, D1556), propidium iodide (Sigma, P4864), FDGlu fluorescent β-glucosidase substrate (Sigma, 92396), Promega FastBreak lysis reagent (V8571), Tween 80 (Sigma, P1754), D-(+)-Trehalose Dihydrate (Thermo Fisher, 182550250), Betaine HCl (Sigma, B3501), PEG 8000 (Hampton Research, HR2-535), FluorN 2900Ni (Cytonix), FITC-dextran (Sigma, FD40S), and 1H,1H,2H,2H-perfluoro-1-octanol (Sigma, 370533).

### Terminology

In this manuscript, microfines refers to small droplets at the low-micron or submicron scale generated during manipulations such as vortexing, homogenization, transfer, or oil exchange. Depending on context, microfines were either an undesired byproduct of droplet generation or were intentionally produced and used to stabilize larger droplets during subsequent handling or thermocycling.

### Preparation of PFPE-PEG reaction product

A PFPE-PEG reaction product was prepared by direct EDC coupling of Krytox 157 FSH to Jeffamine ED900 in a biphasic aqueous/fluorinated system.

Jeffamine ED900 (166.1 mg, 185 µmol) was dissolved in 7.5-10 mL deionized water in a 15 mL conical-bottom tube by vortexing. The solution was buffered to pH 6-7 with MES. Separately, an oil phase was prepared by dissolving 2 g Krytox 157 FSH in 38 g Novec HFE-7500 in a 50 mL conical-bottom tube, corresponding to an approximate 5% (w/w) starting concentration relative to the starting Krytox mass. The Jeffamine:Krytox molar ratio was 0.6, representing a 20% excess of Jeffamine relative to the nominal 0.5 stoichiometric ratio associated with a PFPE-PEG-PFPE architecture.

EDC (300-500 mg) was dissolved in the aqueous phase immediately before use. The aqueous phase was then emulsified into the oil phase by vortexing or shaking. The resulting emulsion was incubated at room temperature on a rocker or rotary shaker for 16-72 h.

### Cleanup of PFPE-PEG reaction product

After incubation, the emulsion was divided between two fresh 50 mL tubes. Dry PEG 35,000 flakes were added to each tube to a depth of approximately 1.5 cm. The tubes were centrifuged at maximum relative centrifugal force (3500-7500xg) for 60 min at room temperature or above.

During centrifugation, aqueous droplets rose and coalesced. Water and water-soluble components were absorbed into the PEG layer, forming a solid or semisolid plug, while the fluorinated phase remained below. The lower fluorinated phase was recovered by passing a syringe or transfer pipette through the PEG layer and decanting into a fresh tube. This cleanup was repeated at least twice until the fluorinated phase no longer had an opaque white appearance. A final water-emulsification cleanup step was then performed to reduce carryover of soluble material. Before use, the recovered fluorinated phase was filtered through a 0.2 µm syringe filter. Working droplet-generation oil was prepared by dilution with fresh oil to 2% surfactant where indicated.

### Preparation of PFPE-Tris reaction product

PFPE-Tris was prepared using the same general procedure, substituting Tris base for Jeffamine ED900. Tris base was added as a 1.1-fold molar excess relative to Krytox 157 FSH, corresponding to 84.7 µL of a 1 M stock solution. Emulsification, incubation, cleanup, and filtration were performed as above.

### Genomic library screening for **β**-glucosidase activity

Genomic DNA was extracted from *Escherichia coli* or *Bacteroides intestinalis*, hydrodynamically sheared using Covaris g-Tubes (Covaris), end repaired, and cloned into pMiniT (New England Biolabs) or pCR-XL-2-TOPO (Invitrogen). Libraries were initially cultivated in DH10β cells (New England Biolabs), after which plasmid libraries were extracted and transformed into NEB T7 Express cells (New England Biolabs).

Expression cultures were induced in mid-log phase with 0.4 mM IPTG. After 4 h of induction, cells were pelleted and resuspended in β-glucosidase assay buffer containing 20% OptiPrep and propidium iodide. Aggregates were removed with a 5 µm syringe filter. This cell suspension was co-flowed with a second aqueous stream containing β-glucosidase buffer, FastBreak lysis reagent, propidium iodide, and FDGlu substrate.

Droplets were generated in a 20 µm × 20 µm flow-focusing device. Oil flow was 10 µL/min, and each aqueous inlet was 1.5-2 µL/min. Droplets were collected into 100 µm-thick capillaries, incubated for 1-3 days. Imaging was performed on a Nikon C2 confocal microscope, using 488 nm and 561 nm lasers to detect hydrolyzed FDGlu (fluorescein) and propidium iodide-stained genetic material, respectively.

### Thermocycling comparison of coupled versus uncoupled systems

A mock PCR aqueous mixture contained 1× KOD polymerase buffer, 2 mM MgSO_4_, 0.2 M trehalose, 1 M betaine, 10 ng/µL polyadenylic acid, 2% Tween 80, 2.5% PEG 8000, and yellow tracking colorant. This mixture was split into two aliquots.

One aliquot was supplemented with 0.75% Jeffamine ED900 and emulsified in HFE-7500 containing 5% (w/w) uncoupled Krytox 157 FSH. The second aliquot was emulsified using 2% PFPE-PEG reaction product in HFE-7500. Both emulsions were generated in a 20 µm × 20 µm flow-focusing in-house dual inlet droplet generator device at 5 µL/min oil and 1 µL/min in each aqueous line using a microfluidic pump (Harvard Apparatus PhD Ultra Syringe Pump 70-3007). Droplets prepared with PFPE-PEG were subsequently transferred into 5% PFPE-PEG in FC-40 for thermocycling.

Emulsions were collected into PCR strip tubes and overlaid with 70 µL mineral oil. Thermocycling consisted of 95°C for 1:00, followed by 35 cycles of 95°C for 0:45, 58°C for 0:47, and 70°C for 0:25. Emulsions were photographed after cycling.

### Microfine-stabilized large-droplet PCR workflow

A 2 mL aqueous microfine mix was prepared containing 1× surfactant-free KOD polymerase buffer, 2 mM MgSO_4_, 0.2 M trehalose, 1 M betaine, 0.01% Tween 80, and 0.05% Cytonix 2900Ni. This mixture was split in half. One half was emulsified into 5% PFPE-PEG in HFE-7500 and the other into Bio-Rad Droplet Generation Oil for EvaGreen. Used oil was decanted and replaced with fresh corresponding droplet generation oil, and the suspensions were homogenized for 2 min at 30,000 rpm to generate microfines.

A 0.4 mL PCR master mix contained 1× surfactant-free KOD buffer, 2 mM MgSO_4_, 400 µM dNTP mix, 0.2 M trehalose, 1 M betaine, 10 ng/µL polyadenylic acid, 0.5 µM each forward and reverse primer, 2.67 X 10copies template, 0.01% Tween 80, 0.05% Cytonix 2900Ni, 62.5 ng/µL BSA, and 12 U KOD polymerase. A 20 µL bulk control reaction was reserved.

The remainder was divided into FITC-dextran-labeled and unlabeled fractions and then subdivided for emulsification with either PFPE-PEG or Bio-Rad droplet-generation oil. Droplets were generated in a 50 µm × 50 µm flow-focusing in-house droplet generator device at 10 µL/min oil and 6 µL/min total aqueous flow. A portion of each FITC-dextran-labeled sample was retained for pre-PCR imaging.

For thermocycling, the 2% droplet-generation oil was replaced with approximately equal volume of corresponding microfines. Samples were placed in PCR tubes, overlaid with 30 µL mineral oil, and cycled using 95°C for 2:00 followed by 35 cycles of 95°C for 0:25, 58°C for 0:15, and 70°C for 0:10.

After thermocycling, unlabeled samples were lysed with 1H,1H,2H,2H-perfluoro-1-octanol and diluted to approximately 10% in TE buffer for qPCR analysis. FITC-dextran-labeled samples were washed with used droplet-generation oil to remove microfines and then imaged in capillaries.

### Protein crystallization in droplets

The sialate O-acetylesterase from *Phocaeicola vulgatus* ATCC 8482 (ABR41743.1) was expressed and purified from *E. coli* as described previously [8,9] and buffer exchanged to crystallization buffer (20 mM HEPES, pH 8.0, 250 mM NaCl) at 78 mg/mL concentration. This sample was co-flowed with 2× MCSG4-B2 screen solution containing 0.1 M Bis-Tris propane, pH 7.0, 1.2 M potassium sodium tartrate, and 1% Cytonix 2900Ni. Droplets were generated in a 50 µm × 50 µm flow-focusing device using either 5% PFPE-PEG reaction product in HFE-7500 or Bio-Rad Droplet Generation Oil for Probes. Flow rates were 5 µL/min oil, 1 µL/min protein solution, and 1 µL/min screen solution.

### Stability trials for problematic protein crystallization screens

Protein crystallization screens previously found to destabilize Bio-Rad Droplet Generation Oil for Probes were retested using 5% PFPE-PEG reaction product in HFE-7500 with 1× aqueous screens containing 1% Cytonix 2900Ni. For each condition, 250 µL oil was combined with 100 µL screen in a 0.5 mL microtube and vortexed to emulsification. Samples retaining a white foamy emulsion layer after 72 h were scored as positive. Samples with very large droplets or visible demulsification were scored as negative.

### Data presentation

Most experiments were performed as method-development or application demonstrations rather than formal replicate-rich comparative studies. Accordingly, results are presented as representative practical outcomes and comparative observations. Unsupported claims of chemical purity or fully defined composition are intentionally avoided.

## Results

### 1. Direct EDC coupling produced functional PFPE-based materials for droplet microfluidics

Direct reaction of Krytox 157 FSH with Jeffamine ED900 or Tris under EDC-mediated conditions yielded fluorinated materials that could be recovered after PEG-assisted cleanup and used as droplet-stabilizing agents. Because the products were not fully structurally characterized, they are described here as PFPE-PEG and PFPE-Tris reaction products rather than as compositionally defined surfactants. Nonetheless, both materials were sufficiently reproducible in practice for repeated use in microfluidic workflows.

### 2. Head-group identity strongly affected droplet-generation behavior

The two principal reaction products differed in droplet behavior. PFPE-Tris at 5% in HFE-7500 was useful for generating relatively uniform small droplets in our devices (Figs. 1, 2, 5, and 6). Under similar conditions, PFPE-PEG at 5% more often generated broader droplet populations and visible microfines. For this reason, PFPE-PEG was generally used at 2% for routine droplet generation (Figs. 3 and 4), while PFPE-Tris was preferred when formation of low-diameter, relatively uniform droplets was desired.

**Figure 1.**
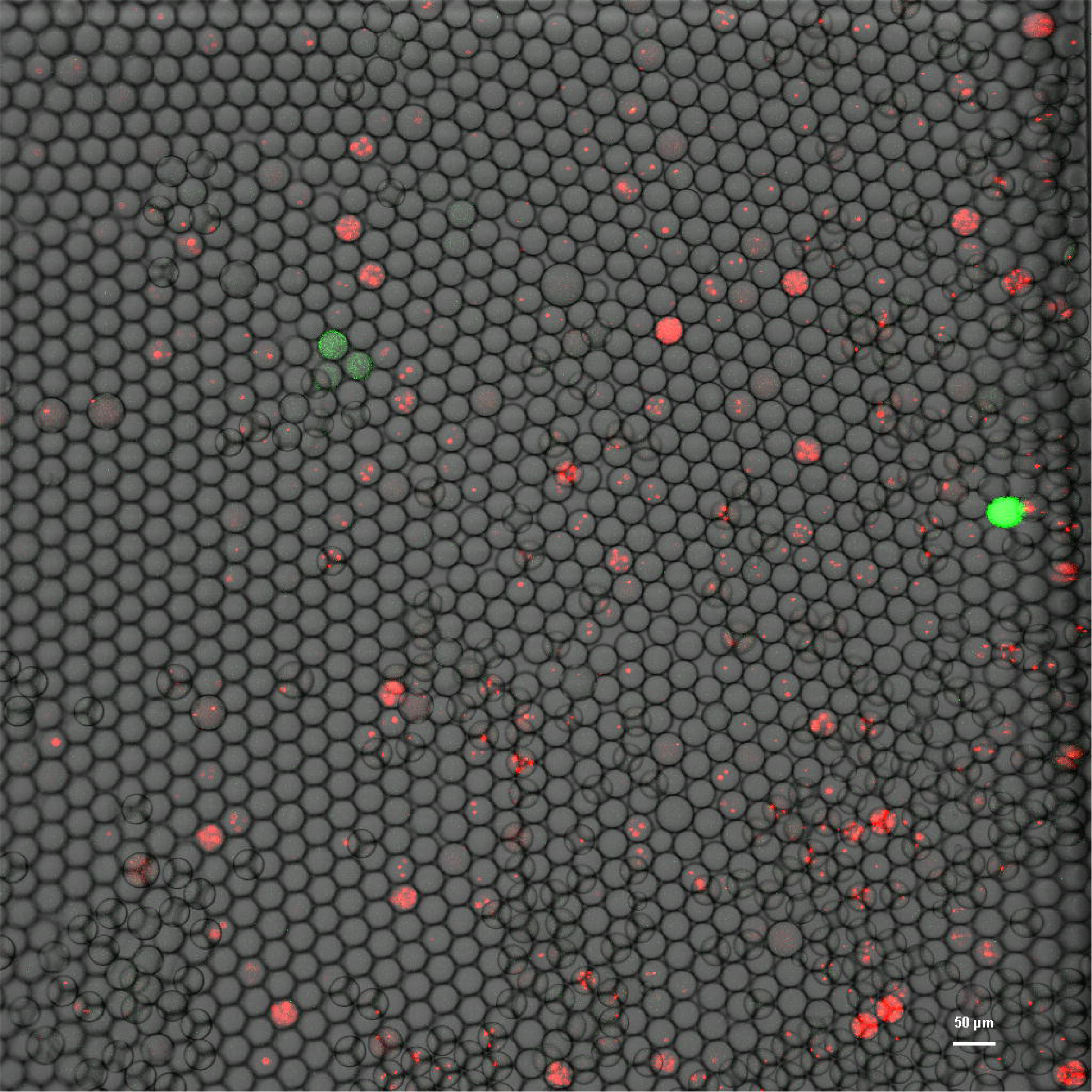
*Bacteroides intestinalis* genomic library screened for β-glucosidase activity in droplets stabilized with 5% PFPE-Tris after 3 d incubation and permeabilized. Green fluorescence indicates FDGlu hydrolysis. Red fluorescence indicates propidium iodide staining of nucleic-acid-containing cellular material.

**Figure 2.**
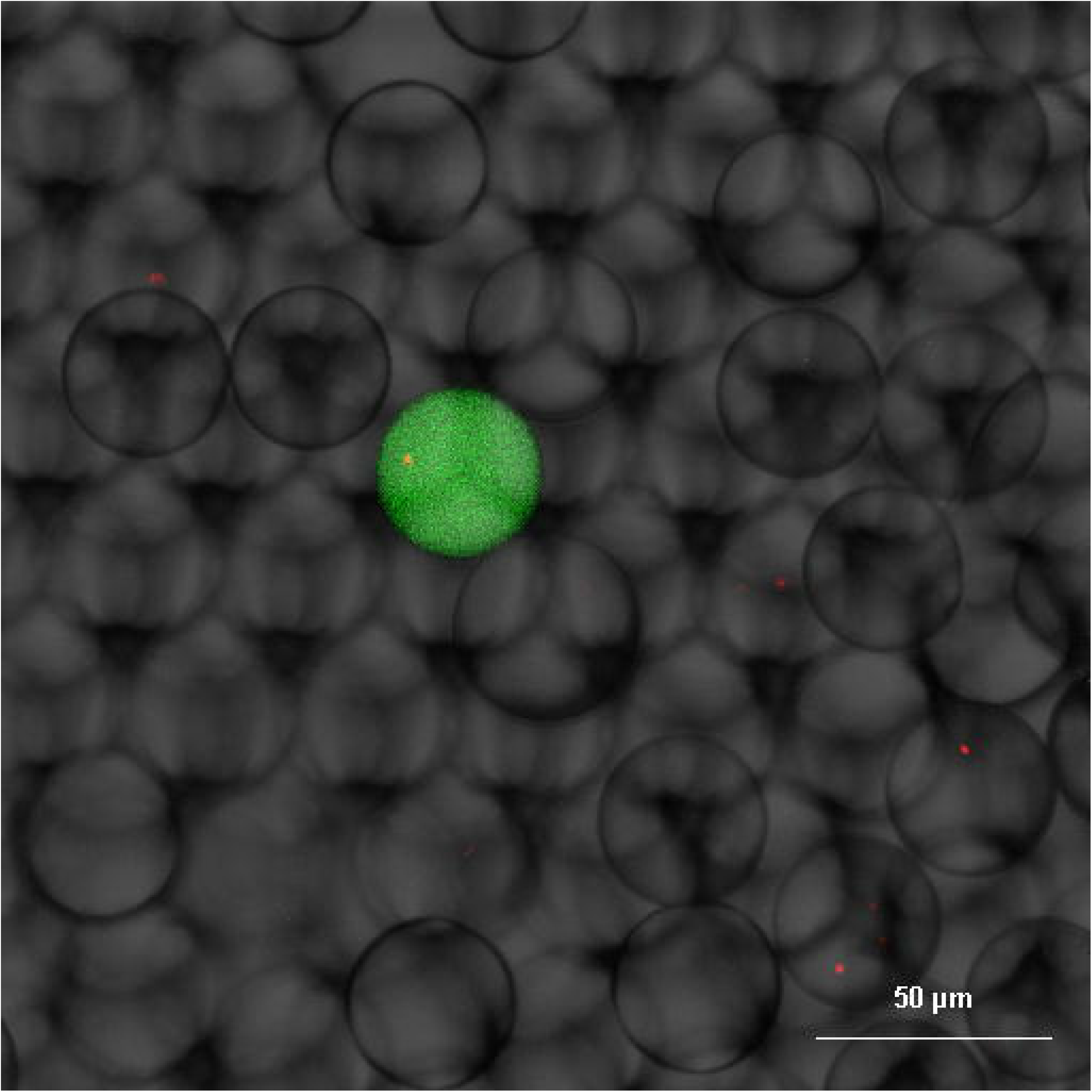
Additional field of view from the *B. intestinalis* β-glucosidase screen shown in Figure 1.

**Figure 3.**
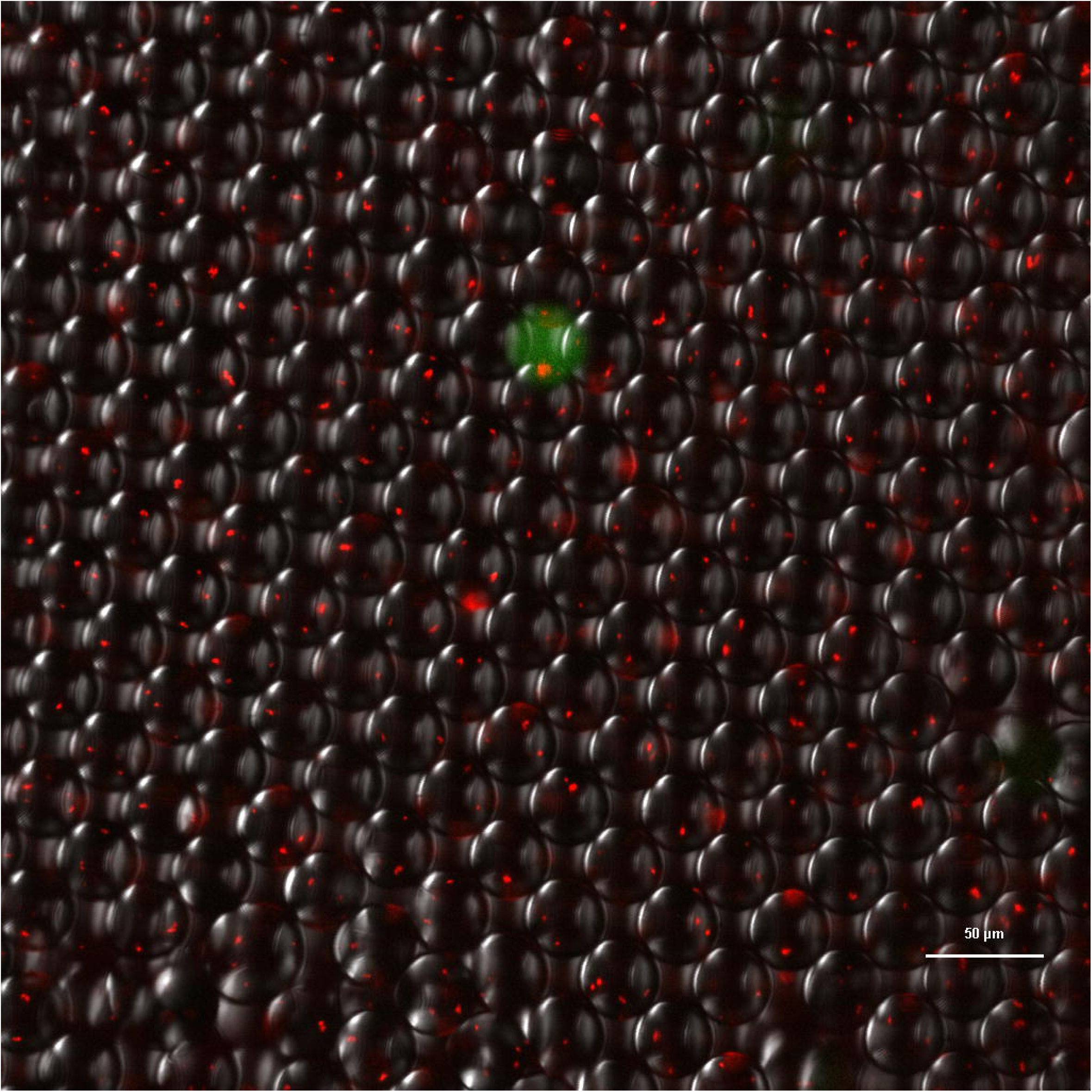
*Escherichia coli* expression library screened for β-glucosidase activity in droplets stabilized with PFPE-PEG.

**Figure 4.**
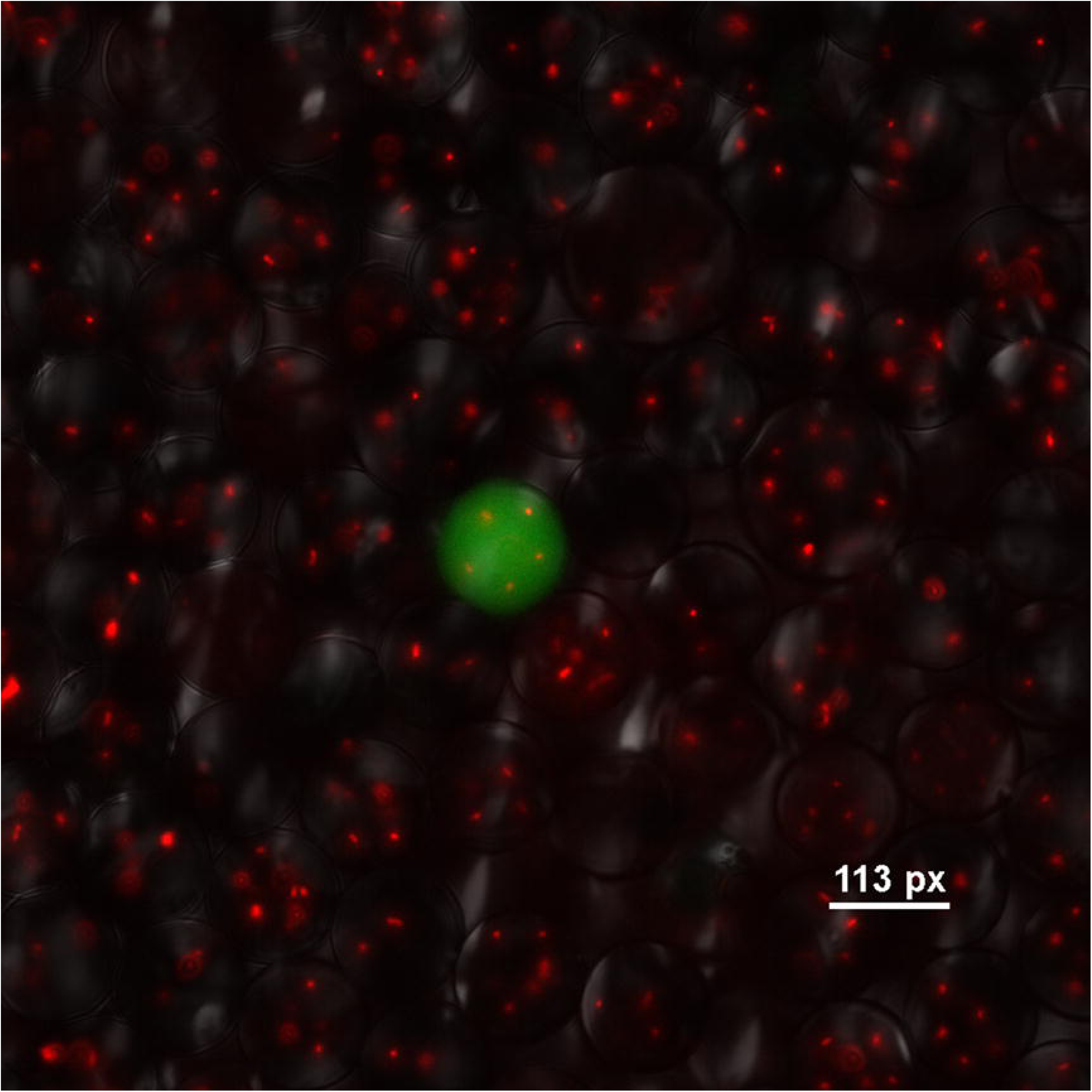
Additional field of view from the *E. coli* β-glucosidase screen shown in Figure 3.

**Figure 5.**
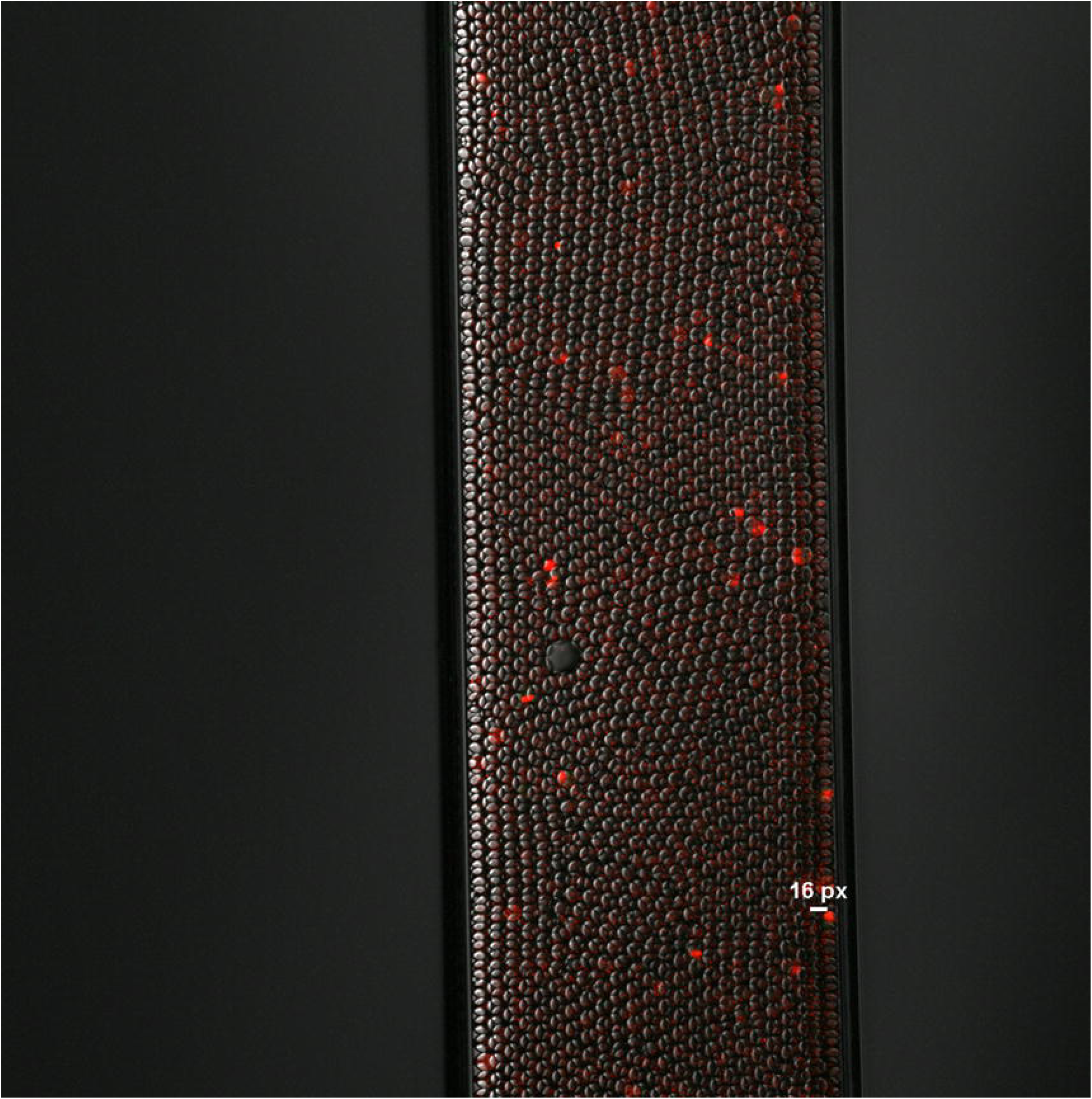
*B. intestinalis* genomic library screened for β-glucosidase activity in droplets stabilized with 5% PFPE-Tris after 1 day incubation.

**Figure 6.**
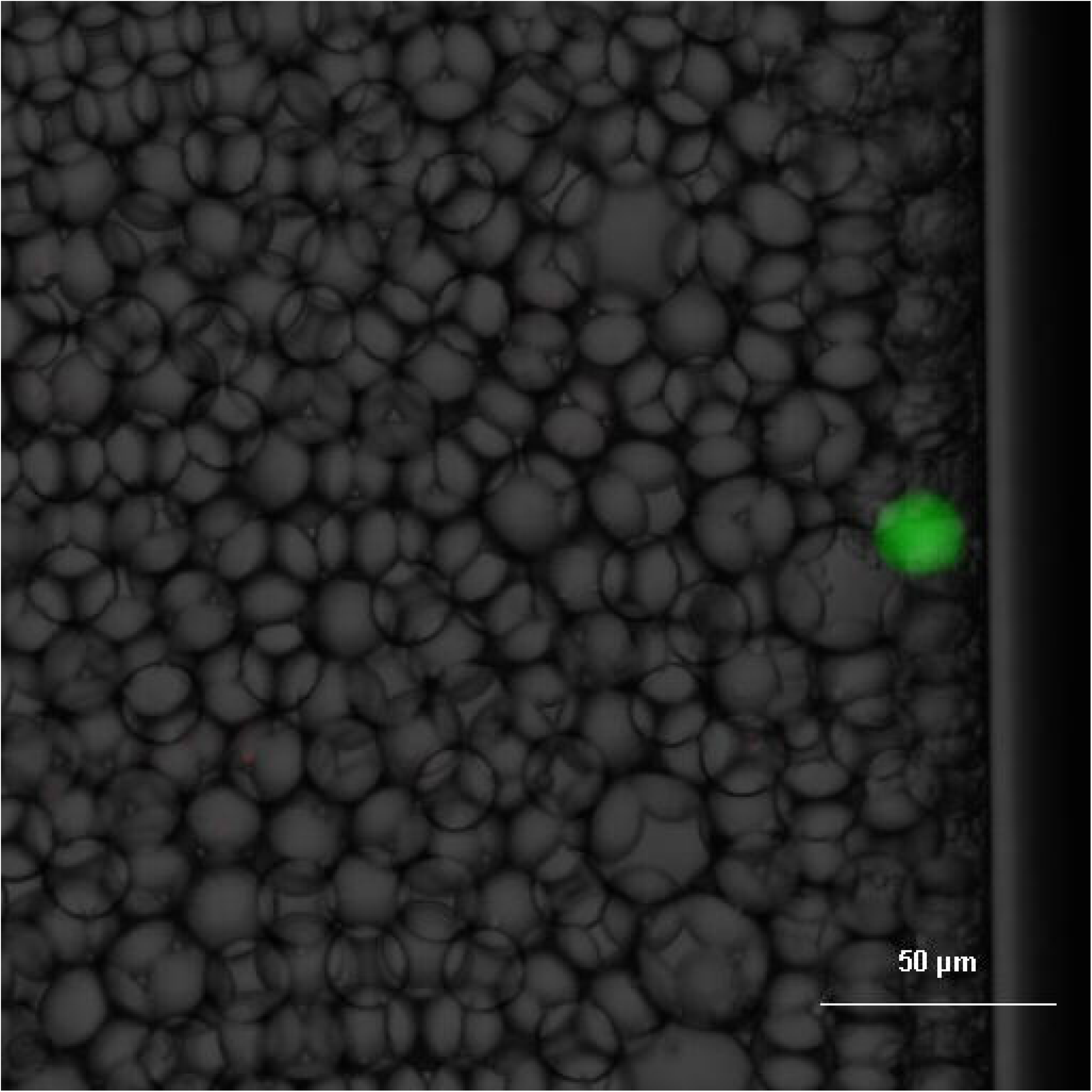
Additional field of view from the *B. intestinalis* screen shown in Figure 5.

### 3. PFPE-based reaction products supported genomic library screening for **β**-glucosidase activity

Both PFPE-Tris and PFPE-PEG were compatible with droplet-based genomic library screening for β-glucosidase activity. In these assays, positive droplets displayed green fluorescence from FDGlu hydrolysis, while propidium iodide provided red fluorescence associated with nucleic-acid-containing material from lysed cells. Positive droplets were observed in screens derived from both *Bacteroides intestinalis* and *E. coli* (Figs. 1-6). Typical droplet diameters were approximately 18-20 µm, and each 100 µm-thick capillary contained approximately four to five droplet layers. These observations indicate that the reaction products supported cell loading, lysis, substrate retention, and assay readout under the conditions tested.

### 4. Coupled PFPE-PEG outperformed an uncoupled Jeffamine/Krytox mixture in a mock thermocycling workflow

In a mock PCR comparison, droplets stabilized with the PFPE-PEG reaction product showed better practical persistence through thermocycling than droplets produced using an uncoupled aqueous Jeffamine ED900 plus fluorinated Krytox 157 FSH approach (Fig. 7). Under our conditions, we did not reproduce comparable stability with the uncoupled system. This result supported continued use of the coupled PFPE-PEG material for temperature-stressed workflows.

**Figure 7.**
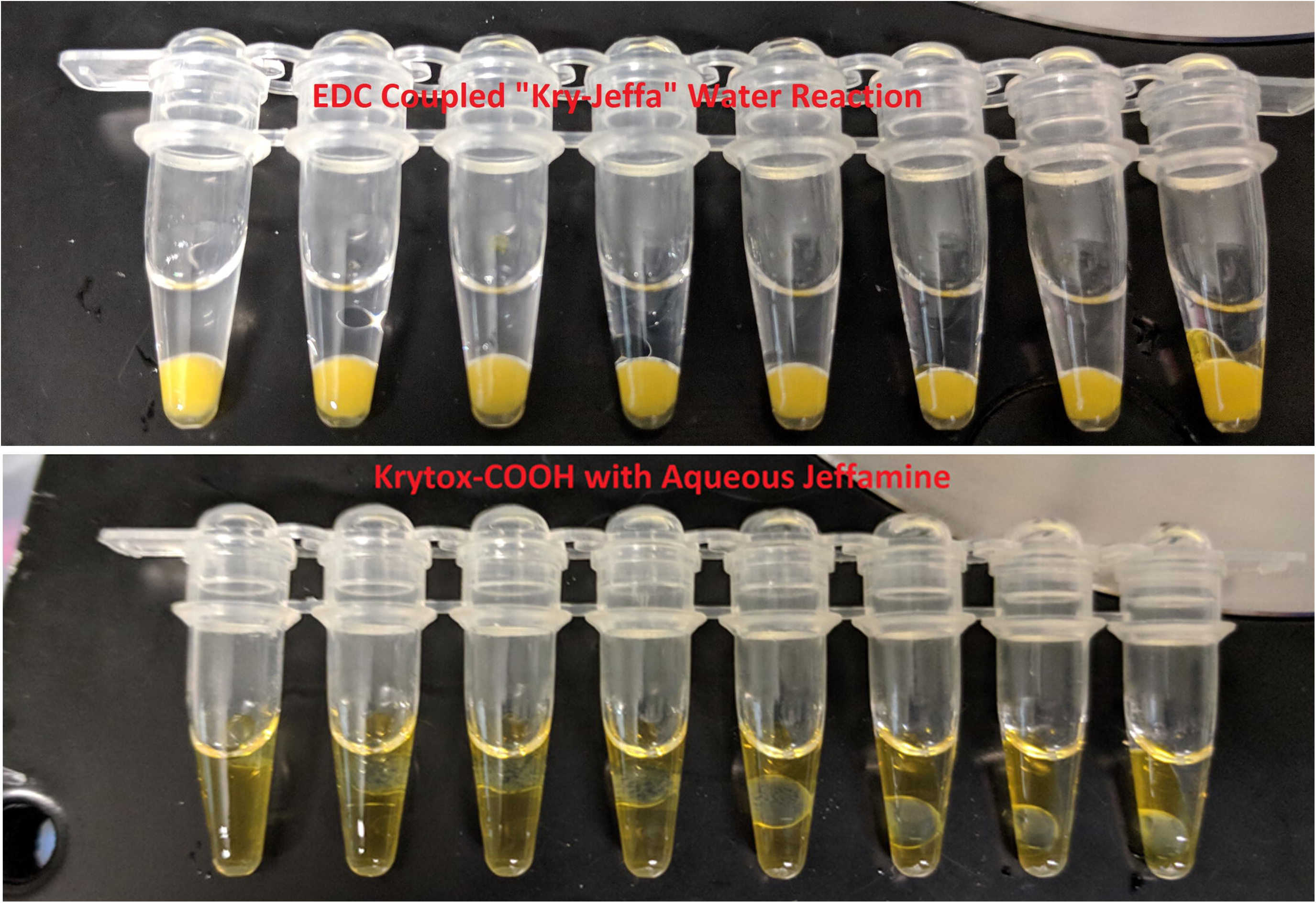
Mock thermocycling comparison of droplets generated with PFPE-PEG reaction product versus an uncoupled Jeffamine ED900/Krytox 157 FSH system. Yellow regions correspond to aqueous material or droplets; the clear upper phase is mineral oil.

### 5. Thermocycling experiments revealed a distinction between apparent emulsion stability and retained compartmentalization

Across several thermocycling-associated workflows, apparent post-PCR emulsion integrity did not necessarily indicate preserved droplet compartmentalization. Small-droplet samples could appear intact by gross inspection and still fail to provide convincing evidence of true in-droplet amplification under our conditions. For larger droplets, microfines were often required to preserve apparent stability during thermocycling, but they also complicated post-cycle assessment by masking droplet failure.

This distinction was illustrated in the comparison between PFPE-PEG and Bio-Rad EvaGreen oil. qPCR of recovered material initially suggested greater amplification-associated signal in the Bio-Rad condition (Fig. 8). However, analysis of corresponding FITC-dextran-labeled samples showed near-complete droplet loss after thermocycling and washing in the Bio-Rad condition (Figs. 9 and 10A), indicating that recovered nucleic acid signal was not sufficient evidence of preserved droplets. In contrast, PFPE-PEG samples retained visible droplets after thermocycling and washing (Fig. 10B), although recovered qPCR signal was lower. These results support the use of direct imaging alongside molecular readouts when evaluating thermocycled droplet assays.

**Figure 8.**
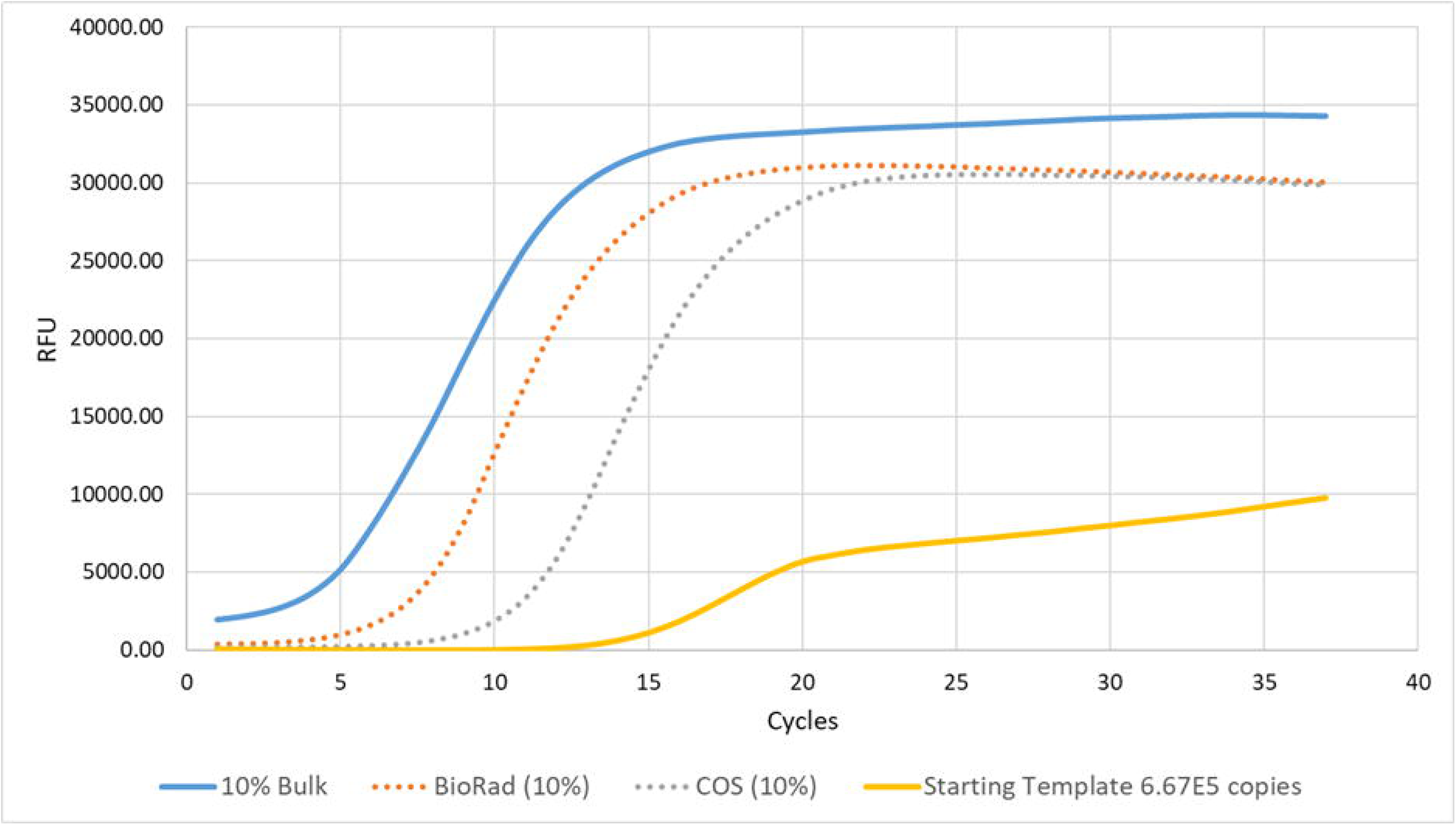
Comparative qPCR of recovered material from microfine-stabilized large-droplet thermocycling experiments using PFPE-PEG or Bio-Rad EvaGreen oil.

**Figure 9.**
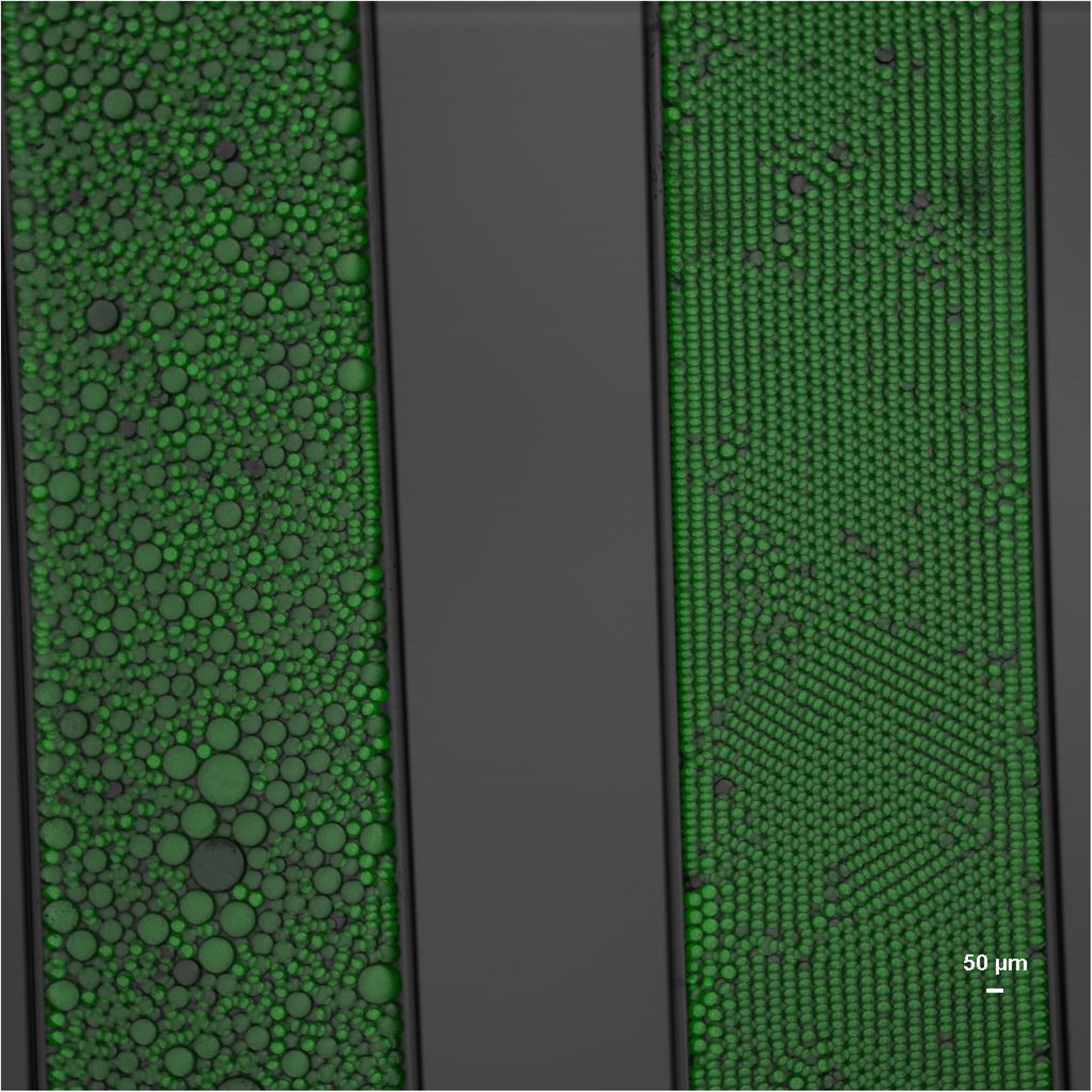
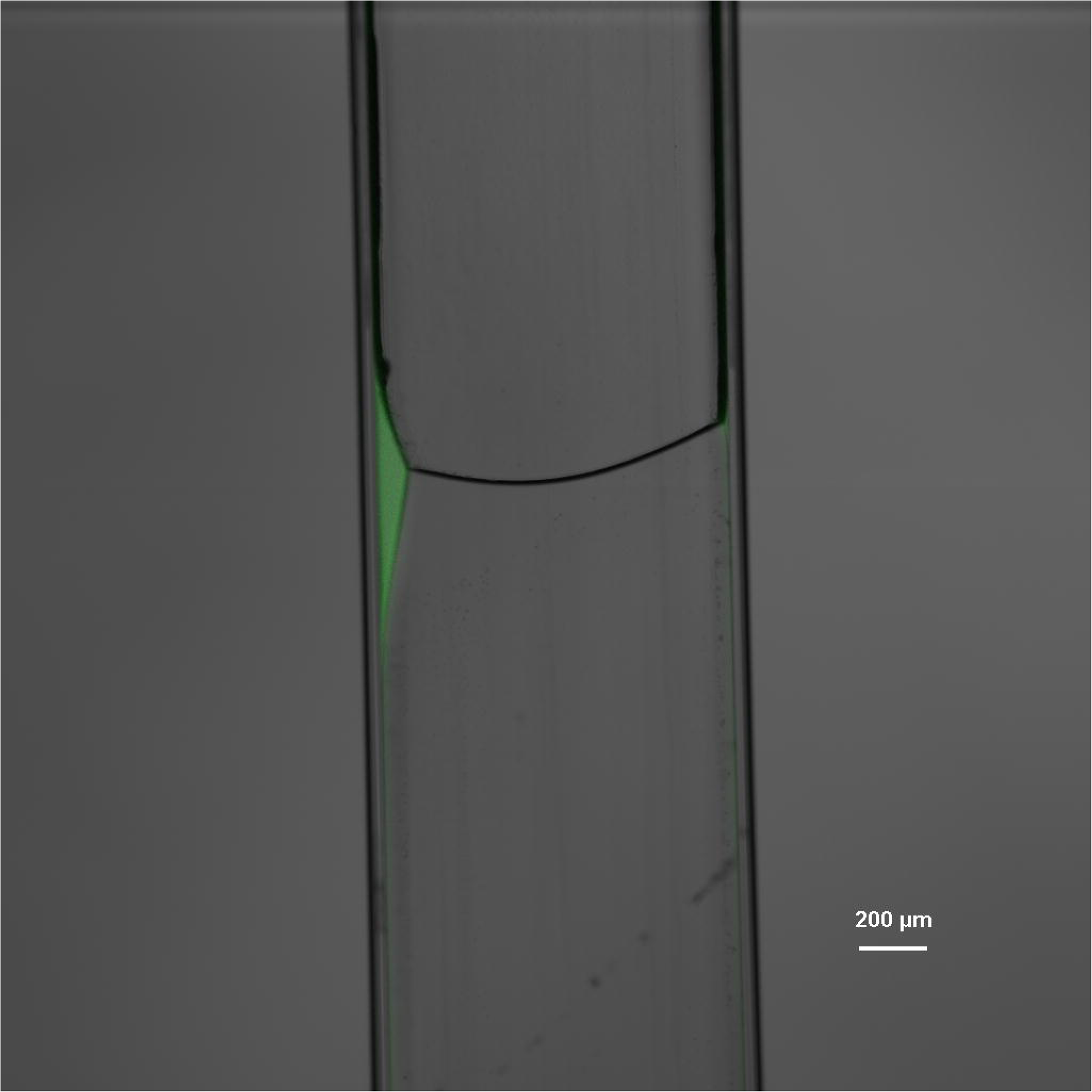
FITC-dextran-labeled droplets before thermocycling.

**Figure 10.**
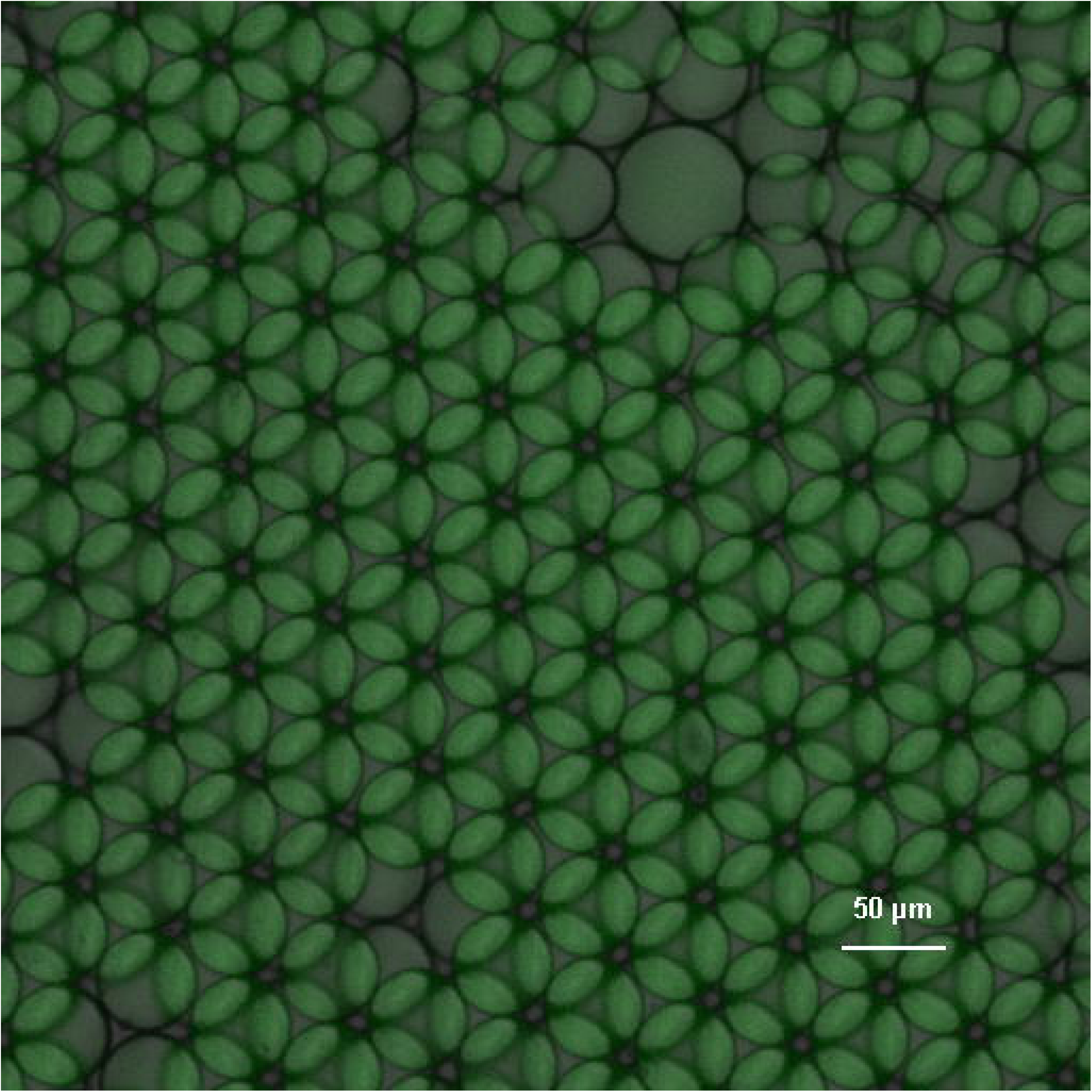
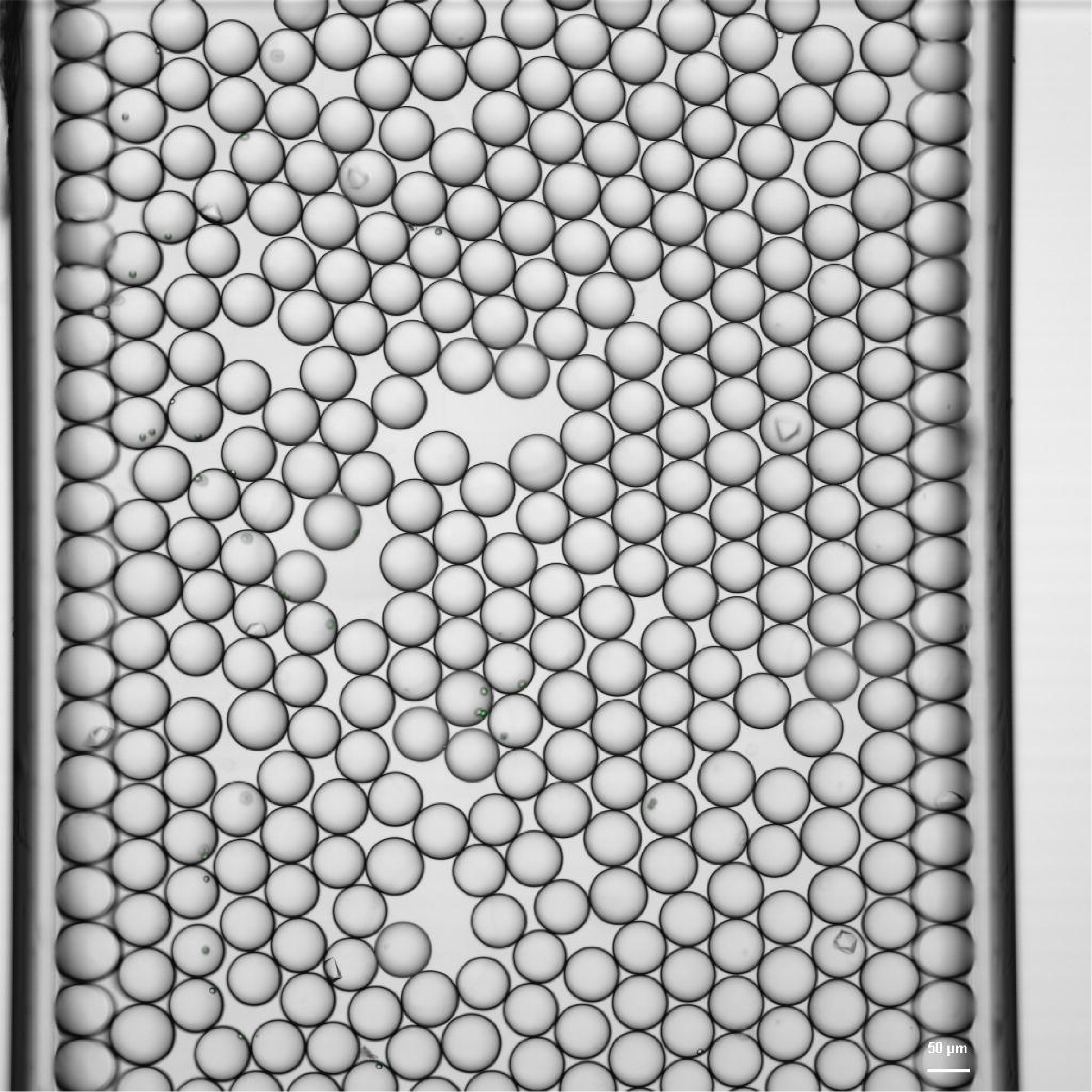
Post-thermocycling FITC-dextran samples after washing to remove microfines. (A) Bio-Rad condition, in which intact droplets were not recovered. (B) PFPE-PEG condition, in which visible droplets remained after cycling and washing.

### 6. PFPE-PEG supported protein crystallization droplets and improved behavior in many problematic screen conditions

The PFPE-PEG reaction product was also compatible with protein crystallization workflows [10]. In a representative crystallization assay, droplets generated with PFPE-PEG in HFE-7500 were visually comparable to droplets generated with Bio-Rad Droplet Generation Oil for Probes (Figs. 11 and 12).

**Figure 11.**
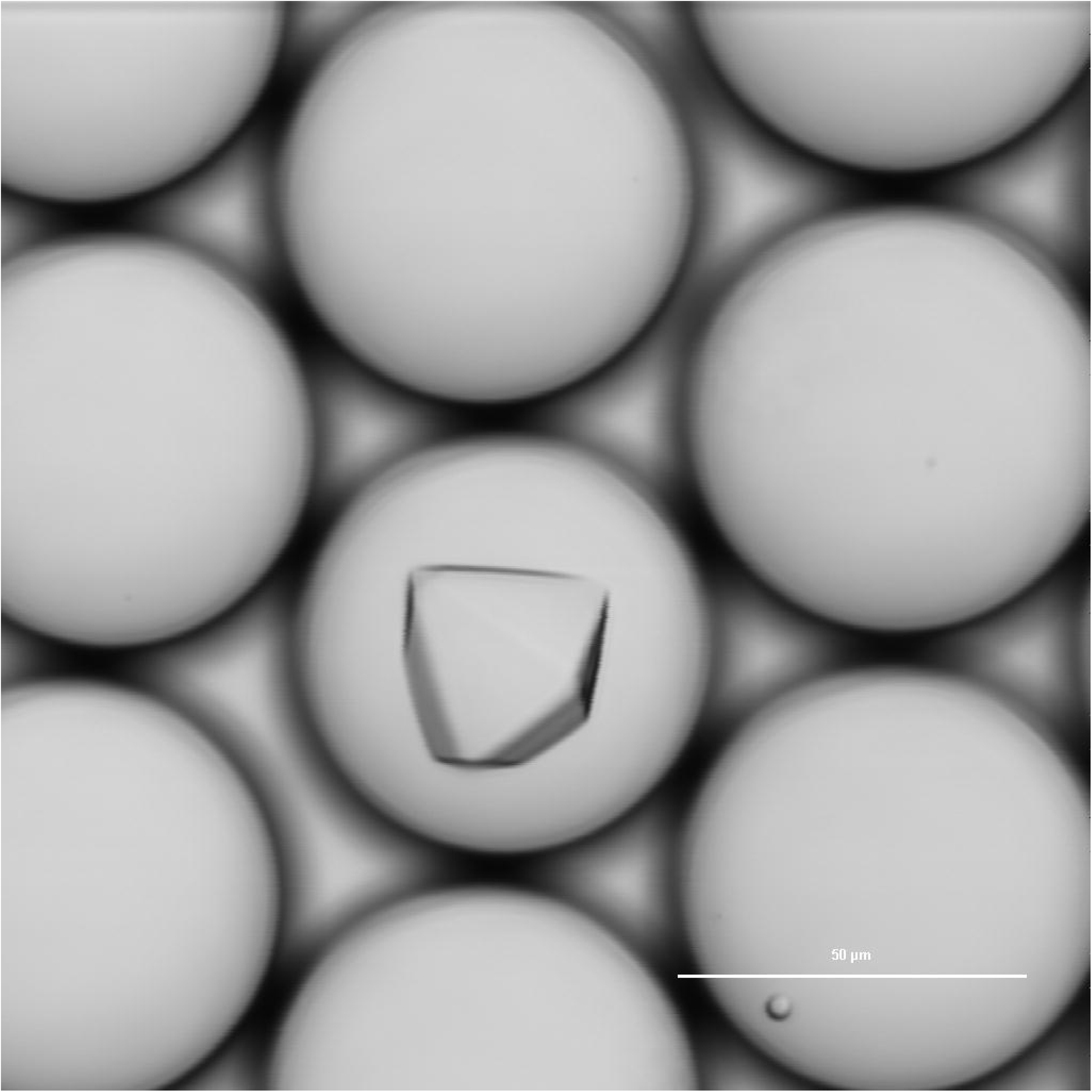
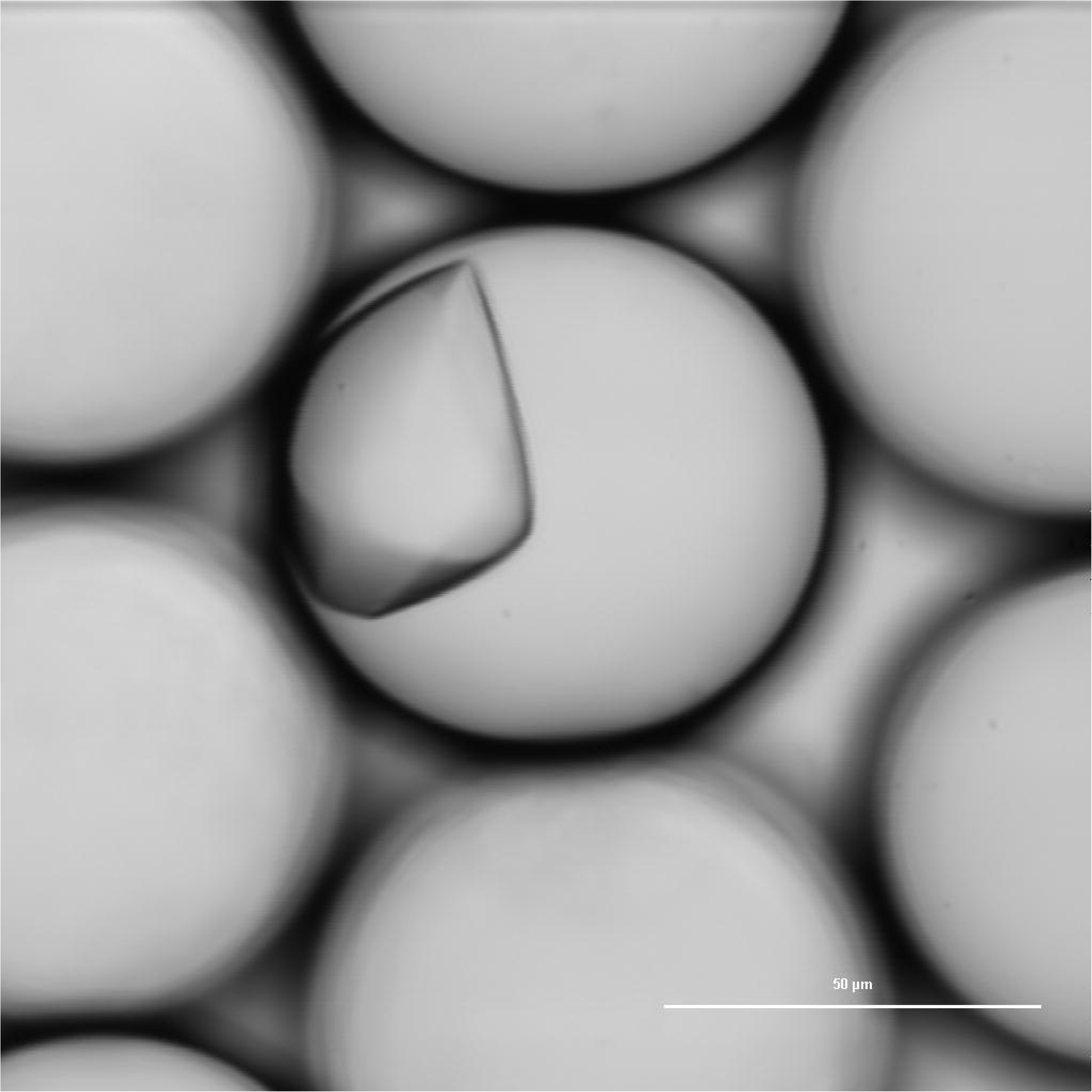
Protein crystallization droplets generated using Bio-Rad Droplet Generation Oil for Probes.

**Figure 12.**
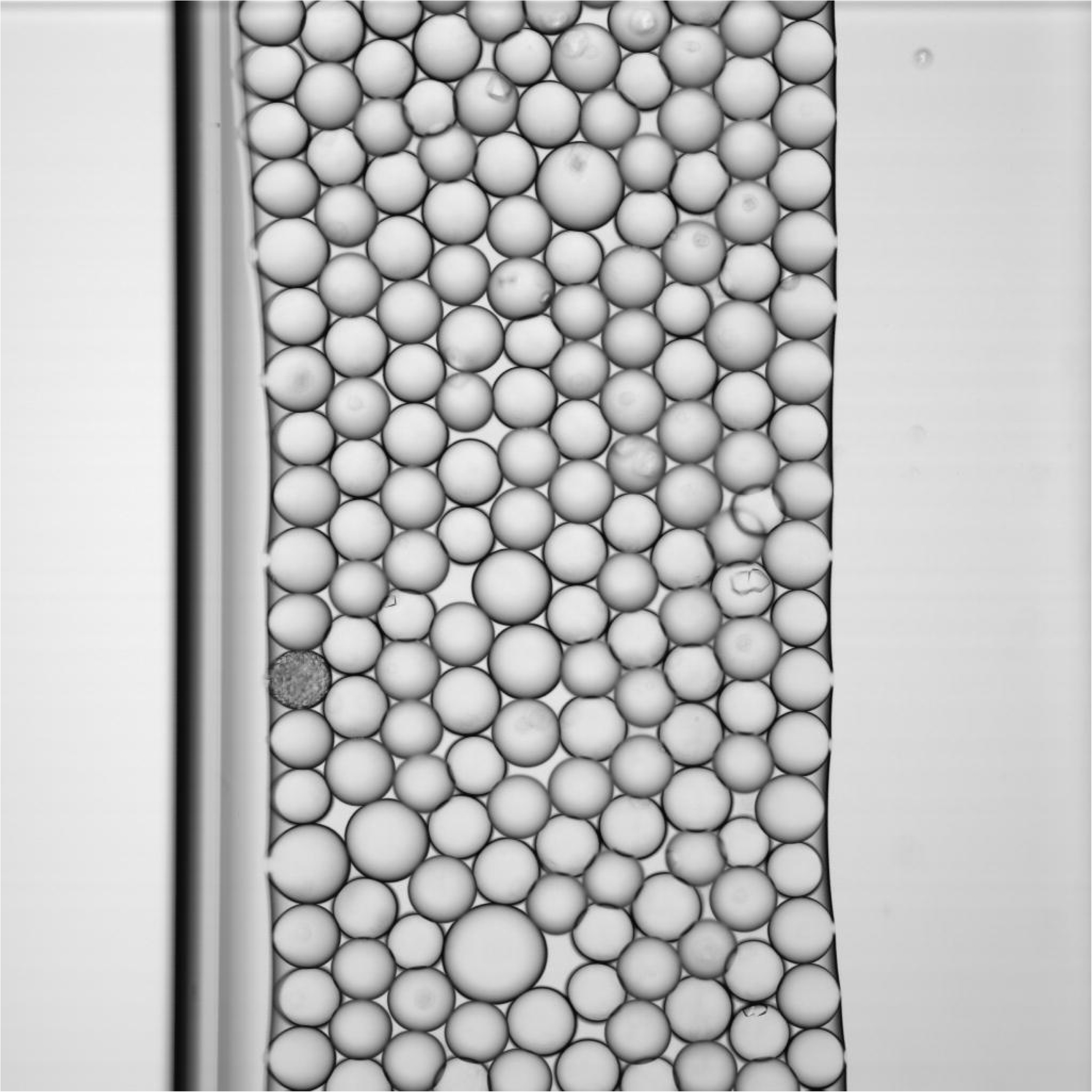
Protein crystallization droplets generated using 5% PFPE-PEG reaction product in HFE-7500.

To assess broader utility, 104 crystallization screen conditions previously found to be unstable in the Bio-Rad oil used in our laboratory were retested with PFPE-PEG. Under the assay conditions used, 95 of these conditions were scored as improved or acceptable with PFPE-PEG (Supplementary Table 1). These results suggest that the PFPE-PEG preparation may be useful in crystallization workflows that challenge some commercial droplet oils.

## Discussion

This work describes a practical route for generating functional PFPE-based fluorinated materials using equipment and methods accessible to biology-oriented laboratories. The principal advantage of the workflow is operational simplicity rather than chemical definition. Direct EDC coupling avoided acid chloride preparation and enabled production of usable fluorinated materials with a vortex mixer, centrifuge, standard tubes, and filtration.

The results also underscore the importance of head-group selection. PFPE-Tris and PFPE-PEG showed different behavior in droplet generation and application range, illustrating that accessible in-house preparation can enable formulation-specific optimization. This flexibility may be useful for groups seeking to tune droplet size, oil compatibility, or assay performance.

Several limitations should be stated clearly. First, the products were not fully structurally characterized, so the precise composition of the recovered fluorinated materials remains unknown. The present conclusions are therefore functional rather than structural. Second, many results are comparative and application-based rather than derived from formal benchmarking with extensive replicates. Third, some observations, especially those involving thermocycled droplets, are best interpreted as practical cautions rather than universal rules for all droplet PCR systems.

One important practical finding was the distinction between visual emulsion persistence and true preservation of droplet compartmentalization after thermocycling. In our hands, molecular readouts from broken emulsions could be misleading if interpreted without post-cycle imaging. This issue is relevant to biomicrofluidic assay design more broadly and may merit more systematic future study.

### Conclusions

Direct carbodiimide coupling of Krytox 157 FSH to amine-containing head groups provides a practical, biology-laboratory-compatible route to functional PFPE-based materials for droplet microfluidics. Under the conditions tested, PFPE-Tris was useful for generating relatively uniform small droplets, and PFPE-PEG supported a broader range of biomicrofluidic applications including β-glucosidase screening and protein crystallization. Although additional analytical characterization is needed to define product composition, the workflow offers a useful and adaptable alternative to exclusive reliance on commercial fluorosurfactants.

## Author contributions

Chase Akins conceived the study, performed experiments, and drafted the manuscript. Jessica Johnson performed experiments. Gyorgy Babnigg analyzed data and drafted the manuscript. All authors revised and approved the final manuscript.

## Acknowledgments

We thank Mike Endres for discussions and technical support. We also acknowledge the Advance Protein Characterization Facility for protein purification and characterization resources.

## Data availability

All data supporting the findings of this study are included in the manuscript and supplementary materials or are available from the corresponding author upon reasonable request.

## Supplementary Information

**Supplementary Table 1.** Recovery of problematic protein crystallization screen conditions using 5% PFPE-PEG reaction product in HFE-7500 compared with Bio-Rad Droplet Generation Oil for Probes.

## References

1. Chiu, Y. L., Chan, H. F., Phua, K. K., Zhang, Y., Juul, S., Knudsen, B. R., Ho, Y. P., and Leong, K. W. Synthesis of fluorosurfactants for emulsion-based biological applications. ACS Nano 8, 3913–3920 (2014). doi:10.1021/nn500810n

2. Lee, M., Collins, J. W., Aubrecht, D. M., Sperling, R. A., Solomon, L., Ha, J. W., Yi, G. R., Weitz, D. A., and Manoharan, V. N. Synchronized reinjection and coalescence of droplets in microfluidics. Lab Chip 14, 509–513 (2014). doi:10.1039/C3LC51214B

3. DeJournette, C. J., Kim, J., Medlen, H., Li, X., Vincent, L. J., and Easley, C. J. Creating biocompatible oil-water interfaces without synthesis: direct interactions between primary amines and carboxylated perfluorocarbon surfactants. Anal. Chem. 85, 10556–10564 (2013). doi:10.1021/ac4026048

4. Zeng, Y., Novak, R., Shuga, J., Smith, M. T., and Mathies, R. A. High-performance single cell genetic analysis using microfluidic emulsion generator arrays. Anal. Chem. 82, 3183–3190 (2010). doi:10.1021/ac902683t

5. Holtze, C., Rowat, A. C., Agresti, J. J., Hutchison, J. B., Angilè, F. E., Schmitz, C. H., Köster, S., Duan, H., Humphry, K. J., Scanga, R. A., Johnson, J. S., Pisignano, D., and Weitz, D. A. Biocompatible surfactants for water-in-fluorocarbon emulsions. Lab Chip 8, 1632–1639 (2008). doi:10.1039/B806706F

6. Pan, M., Lyu, F., and Tang, S. K. Fluorinated Pickering emulsions with nonadsorbing interfaces for droplet-based enzymatic assays. Anal. Chem. 87, 7938–7943 (2015). doi:10.1021/acs.analchem.5b01753

7. Chowdhury, M. S., Zheng, W., Kumari, S., Heyman, J., Zhang, X., Dey, P., Weitz, D. A., and Haag, R. Dendronized fluorosurfactant for highly stable water-in-fluorinated oil emulsions with minimal inter-droplet transfer of small molecules. Nat. Commun. 10, 4546 (2019). doi:10.1038/s41467-019-12462-5

8. Babnigg G, Sherrell D, Kim Y, Johnson JL, Nocek B, Tan K, Axford D, Li H, Bigelow L, Welk L, Endres M, Owen RL, Joachimiak A. 2022. Data collection from crystals grown in microfluidic droplets. Acta Crystallographica Section D: Structural Biology 78:997–1009. doi:10.1107/S2059798322004661

9. Sherrell DA, Lavens A, Wilamowski M, Kim Y, Chard R, Lazarski K, Rosenbaum G, Vescovi R, Johnson JL, Akins C, Chang C, Michalska K, Babnigg G, Foster I, Joachimiak A. 2022. Fixed-target serial crystallography at the Structural Biology Center. Journal of Synchrotron Radiation 29:1141–1151. doi:10.1107/S1600577522007895

10. Kim Y, Babnigg G, Jedrzejczak R, Eschenfeldt WH, Li H, Maltseva N, Hatzos-Skintges C, Gu M, Makowska-Grzyska M, Wu R, An H, Chhor G, Joachimiak A. 2011. High-throughput protein purification and quality assessment for crystallization. Methods 55:12–28. doi:10.1016/j.ymeth.2011.07.010

